# Presynaptic mechanism of epileptiform activities in forward-programmed human excitatory neuronal networks

**DOI:** 10.64898/2026.08.04.742761

**Authors:** Jianbin Wen, Jiaqing Li, Michael Peitz, Oliver Brüstle

## Abstract

Epilepsy is one of the most common neurological disorders, yet the mechanisms controlling seizure termination remain poorly understood. In particular, why rhythmic spike-wave discharges decelerate before stopping is unexplained. Here, using human iPSC-derived excitatory neurons differentiated via targeted forward-programming, we report a similar deceleration phenomenon in cultured neuronal networks. These networks exhibit glutamate-dependent, epileptiform ’super-bursts’ with a slowing rhythm from ∼4 Hz to ∼2 Hz. Combining *in silico* simulations and *in vitro* experiments, we correlate this activity pattern with the hierarchical organization of presynaptic vesicle pools. Nested bursts link to the recycling pool (RP), and sub-bursts associate with the readily releasable pool (RRP). Decelerating RP-to-RRP vesicle translocation shortened the super-bursts, indicating that epileptiform dynamics depend heavily on this translocation process. These findings depict human neuronal networks derived from forward-programmed cells as a model for epileptology, revealing a presynaptic framework for rhythmic discharges in excitatory networks.

**Highlights:**

- Human iPSC-derived glutamatergic networks exhibit epileptiform super-bursts
- Super-burst dynamics are governed by a two-pool presynaptic vesicle hierarchy
- Cytochalasin-D disrupts RP-to-RRP translocation and attenuates super-bursts
- Excitatory networks show intrinsic tonic-clonic bi-stability via RRP dynamics

**eTOC blurb:** Brüstle and colleagues use forward-programmed human iPSC-derived glutamatergic networks to model epileptiform activity. Combining multi-electrode array recordings with computational simulations, they demonstrate that epileptiform super-burst dynamics are governed by a hierarchical two-pool presynaptic vesicle system, and reveal an intrinsic tonic-clonic bi-stability in excitatory networks driven by RRP recovery kinetics.

## Introduction

Epileptic seizures are pathologically excessive or synchronous neuronal activity in the brain (Moshé et al., 2015). Rhythmic activity manifests in diverse seizure types, with slowing low frequency activity frequently preceding seizure termination. For example, the termination of tonic-clonic seizures is typically initiated by clonic bursts with a slowing 4-2 Hz rhythm (Bauer et al., 2017; Boido et al., 2014). Decelerating 4-3 Hz rhythmic spike and wave activities can also be seen in adolescent absence seizures (Bai et al., 2010). Deciphering the mechanisms underlying these seizure-terminating rhythmic patterns might not only provide a better understanding of seizures but also potential targets for seizure control.

Stem cell-based in vitro models offer a powerful platform for investigating fundamental aspects of neural development and neurophysiology (Tao and Zhang, 2016; Vadodaria et al., 2020), including epilepsy (Parent and Anderson, 2015). In this context, forward-programming of human pluripotent stem cells (hPSCs) via overexpression of transcription factors such as NGN2 has evolved as a robust method to efficiently generate enriched populations of forebrain glutamate-releasing neurons (iGlutNs) (Zhang et al., 2013). When co-cultured with astrocytes, these forward-programmed neurons develop mature membrane properties and abundant, functional synapses (Rhee et al., 2019). They readily give rise to standardized neuronal networks generating synchronized network bursts (Wen et al., 2022), which makes them an attractive model for exploring epileptiform dynamics and drug screening (Alaverdian et al., 2023; Mao et al., 2024; Zhao et al., 2024).

To complement these biological platforms, *in silico* computational network models are frequently utilized to isolate and mechanistically dissect the variables governing seizure dynamics (Stefanescu et al., 2012). By mathematically manipulating specific physiological parameters, these models can predict how specific variables could shape the dynamics of seizures (Krishnan and Bazhenov, 2011; Lepeu et al., 2024; Liou et al., 2020; Proix et al., 2014). Combining defined cultured neuronal networks with such computational simulations therefore presents a distinct advantage in epileptological studies.

In the present study, we used *in vitro* human glutamatergic neuronal networks generated from human pluripotent stem cells (hPSCs) to explore the neurophysiological mechanism of epileptiform activities. We observed a rhythmic nested bursting activity that displays electrophysiological features of epileptic seizures. Further exploration revealed a critical role of the hierarchical organization of presynaptic vesicle pools in orchestrating this epileptiform activity. Based on results from *in vitro* experimentation and complementary *in silico* simulations, we propose a novel presynaptic framework for epileptiform activity in disinhibited neural networks.

## Results

### Glutamate-dependent epileptiform activities generated in forward-programmed excitatory human neuronal networks

To generate human forebrain neurons, we used previously established hPSC lines carrying a doxycycline inducible human NGN2 expression cassette in both alleles of the AAVS1 “genomic safe harbor” locus (Wen et al., 2023). Overexpression of NGN2 promotes the differentiation of hPSCs into highly enriched forebrain glutamate-releasing neurons (iGlutNs, Supplementary Fig. S1) (Zhang et al., 2013). Co-culturing these neurons with mouse astrocytes promotes their morphological and functional maturation, manifested by increasing dendritic complexity, mature membrane properties and abundant functional synapses with robust short-term plasticity (Peitz et al., 2020; Rhee et al., 2019). Monitoring the electrophysiological activity of these cultures using multi-electrode arrays (Fig. 1A) yielded a characteristic trajectory of neural network development *in vitro*, beginning with sparse single spikes before 2 weeks and progressing to synchronized activities and network bursts (NBs) between 3 and 6 weeks (Wagenaar et al., 2006; Wen et al., 2022). Interestingly, with further maturation, these NBs progressively developed into higher-order nested bursting patterns we designated as “network super-bursts” (NSB). While immature NSB-like patterns could already be observed at earlier stages, unequivocal NSBs became evident after 7 weeks in culture (Fig. 1B, Supplementary Fig. S2A). NSBs typically start with a strong (measured by the enclosed spike count) and less defined NB, followed by a train of weaker but more uniform NBs with increasing intervals from ∼250 ms to ∼600 ms, lasting for 5-20 seconds in total. The increase of inter-sub-burst intervals with increasing number of sub-bursts followed a linear trajectory in neuronal networks derived from 3 different cell lines (iPSC line C14 and hPSC lines H1 and H9) (Fig. 1C). After NSB initiation, sub-burst strength, defined by the number of enclosed spikes, remained stable (Fig. 1D). Some variations of this pattern were also discernible. For instance, in one cell line, NSBs initiated with a sustained ‘tonic’ state and subsequently transitioned into rhythmic ‘clonic’ bursts (Supplementary Fig. S2B); occasional NSBs exceeding 30 seconds were also observed (Supplementary Fig. S2C). Application of glutamate receptor antagonists CNQX and AP5, as well as GABA, could abolish NSBs, whereas the strength of NSBs (measured by the number of the enclosed sub-bursts) remained unaffected by the GABA_A_ and GABA_B_ receptor antagonists bicuculline and CGP35348, respectively (*F* (2, 149) = 0.64, *p*=0.53, Fig. 1E&F). These observations align with the finding that NGN2 overexpression in hPSCs results in highly enriched glutamatergic populations (Meijer et al., 2019). Of note, NSBs could also be observed in bicuculline-treated excitatory-inhibitory mixed cultures (Supplementary Fig. S2D).

**Figure 1.**
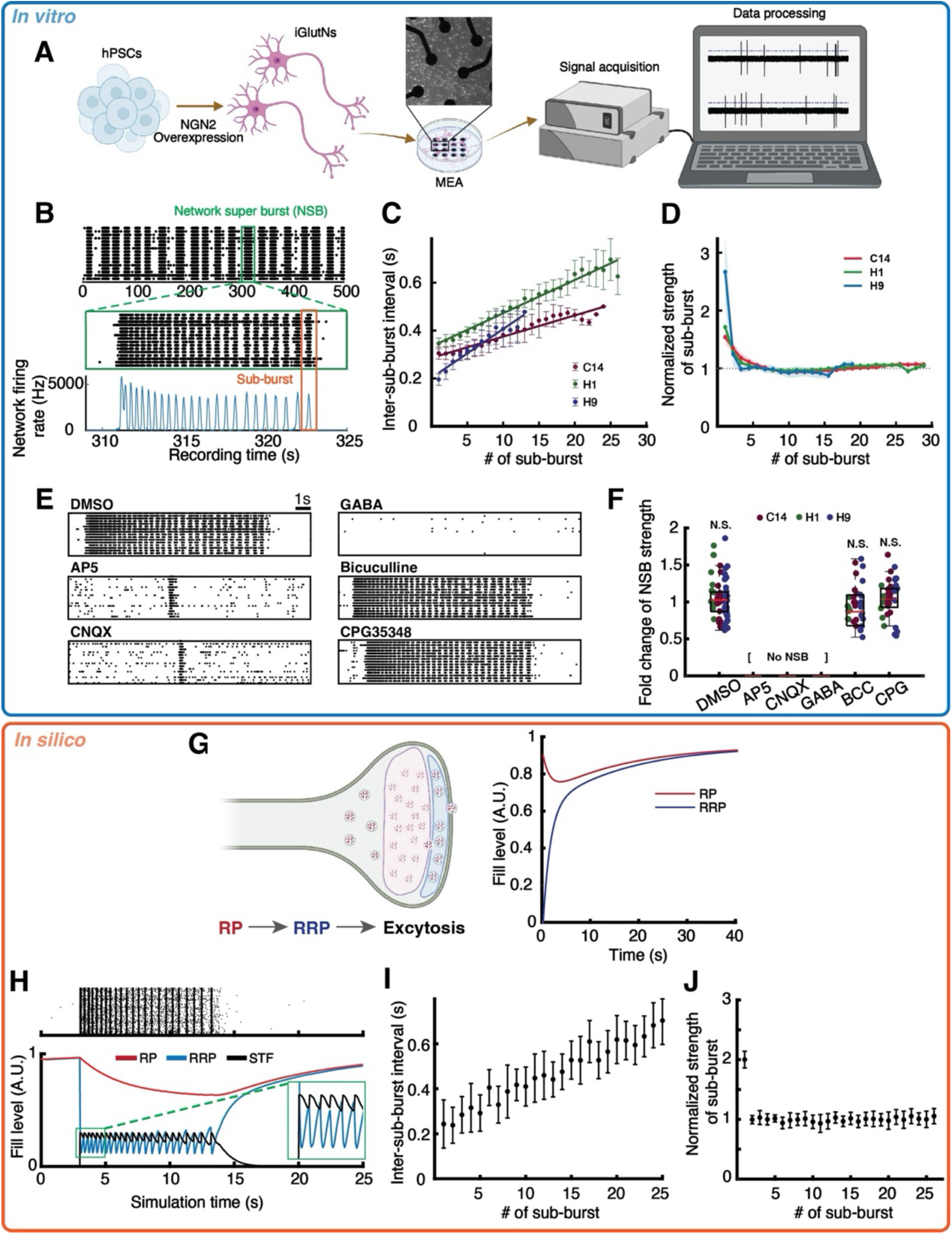
Epileptiform network super-bursts (NSBs) in human glutamatergic neuronal cultures and simulated excitatory neural networks. (**A**) Experimental design. (**B**) Exemplar MEA recording from a 7-week-old culture. One network super-burst (NSB, green box) is enlarged with the corresponding population-level instantaneous network firing rate. Orange box highlights an individual sub-burst (NB) within the NSB. (**C**) Inter-sub-burst intervals (IBIs) within NSBs (linear fits, ≥7 cultures for each background). (**D**) Sub-burst strengths (measured by enclosed spike count) from the recordings in (C), normalized to the median number of spikes in sub-bursts during a NSB. (**E**) Example raster plots showing the effects of AP5 (50μM) and CNQX (10μM), GABA (10μM), bicuculline (10μM) and CGP35348 (1μM) in 7-week-old cultures. (**F**) Change of NSB strength (defined by the number of enclosed sub-bursts) in iGlutN cultures treated with synaptic modulators shown in (E) (≥6 independent MEA cultures per genetic background). N.S., not significant. (**G**) Schematic of sequential translocation of presynaptic vesicles from the recycling pool (RP) to readily-releasable pool (RRP) before exocytosis, and the recovery curves of RP and RRP after RRP depletion. (**H**) Exemplar NSB produced in a simulated excitatory neuronal network model. Dynamics of average RP, RRP and short-term facilitation (STF) in each synapse are shown underneath (RP and RRP normalized to 1, STF adjusted to show crossings with the RRP). (**I-J**) Change of inter-sub-burst intervals (I) and normalized sub-burst strengths (J) within NSBs. Data collected from 20 simulations.

Interestingly, slowing rhythmic bursting activities with a similar frequency range can be seen in adolescent absence seizures (Bai et al., 2010), and at the clonic stage of tonic-clonic seizures (Bauer et al., 2017; Boido et al., 2014). Furthermore, during the clonic phase of tonic-clonic seizures, the spike count in each burst remains stable, but the inter-burst intervals (IBIs) increase steadily from ∼ 250 ms to longer than 1000 ms (Bauer et al., 2017; Boido et al., 2014). These parallels could imply that NSBs in iGlutN cultures share a common electrophysiological mechanism with epileptic seizures. While inhibitory neurons are widely postulated to play pivotal roles in the initiation (and cessation) of seizures (Curtis and Avoli, 2015; Moshé et al., 2015), the here observed emergence of NSBs in networks of limited inhibition suggests an alternative explanation where pure excitatory neurons may suffice to generate epileptiform activities.

### In silico modeling of epileptiform activity patterns

To unravel the critical role of excitatory glutamatergic synapses in the generation of NSB, we adapted computational network models of randomly connected homogeneous excitatory neurons with a single exponential presynaptic dynamics, which showed fast synaptic glutamate depletion and recovery determines the formation of NBs (Loebel and Tsodyks, 2002; Wen et al., 2022). We first modified the in silico model by introducing a synaptic short-term facilitation (STF) to account for the succession of NBs, which enabled the generation of NSBs also featuring a stronger leading NB. However, in contrast to our in vitro observations, the sequence of NBs was unceasing and exhibited constant IBIs (Supplementary Fig. S3). This suggests that STF alone is insufficient, and a slower variable is needed for increasing sub-burst intervals. Along this line, Jirsa et al. previously showed that a minimal dynamical model of seizure activity requires a slow ’permittivity’ variable, in addition to the faster variables governing individual discharges, to account for the waxing and waning time-course of epileptiform activity (Jirsa et al., 2014). One such candidate parameter arises from the concept of a hierarchy of presynaptic vesicle pools: transmitter-containing vesicles must traverse a larger slow-recovering recycling pool (RP) and then the readily releasable pool (RRP) before exocytosis. And upon depletion of RRP, RP would first decrease to refill the RRP, then recover with a slow time constant. Meanwhile, the RRP would first refill rapidly and then be limited by the slow recovery of RP, resulting in a double-exponential recovery process with time constants of <1s and ∼10s, respectively (Rizzoli and Betz, 2005) (Fig. 1G). We therefore incorporated the two-pool presynaptic vesicle dynamics into the computational model. This successfully recapitulated NSBs *in silico*, which had a strong leading NB, followed by homogenous NBs with increasing IBIs – and eventual termination (Fig. 1, H-J). Furthermore, the model elucidated how RP, RRP and STF could interactively drive the network activity during a NSB (Fig. 1H). First, a spontaneous or evoked NB depletes the RRP, silencing the network but also inducing a STF, which promotes another NB after the RRP recovers to a presumed threshold level. The depletion/recovery cycles of the RRP then gradually consume the RP, which in turn slows down the replenishment of the RRP, thereby increasing the IBIs. A bifurcation point occurs when the RRP recovers too slowly to reach the threshold before the STF wanes, leading to the termination of NSBs. After that, the network returns to a resting state as RP and RRP recover.

### Epileptiform activities are associated with presynaptic vesicle pool dynamics

The observations from the *in silico* model would predict that (i) an external stimulus applied shortly following the termination of a NSB and prior to the full recovery of the presynaptic RP should be capable of inducing another, though significantly weaker, NSB, and (ii) the change in NSB strength across increasing recovery time should mirror the recovery curve of the presynaptic RP. Indeed, this was observed *in silico*. Networks stimulated at 10 s intervals showed weaker NSBs (Fig. 2A_i_), whereas stimulation at 20 s intervals yielded stronger NSBs (Fig. 2A_ii_). The recovery of NSBs could be fitted to an exponential curve with a time constant close to that preset for RP recovery in the model (Fig. 2B).

**Figure 2.**
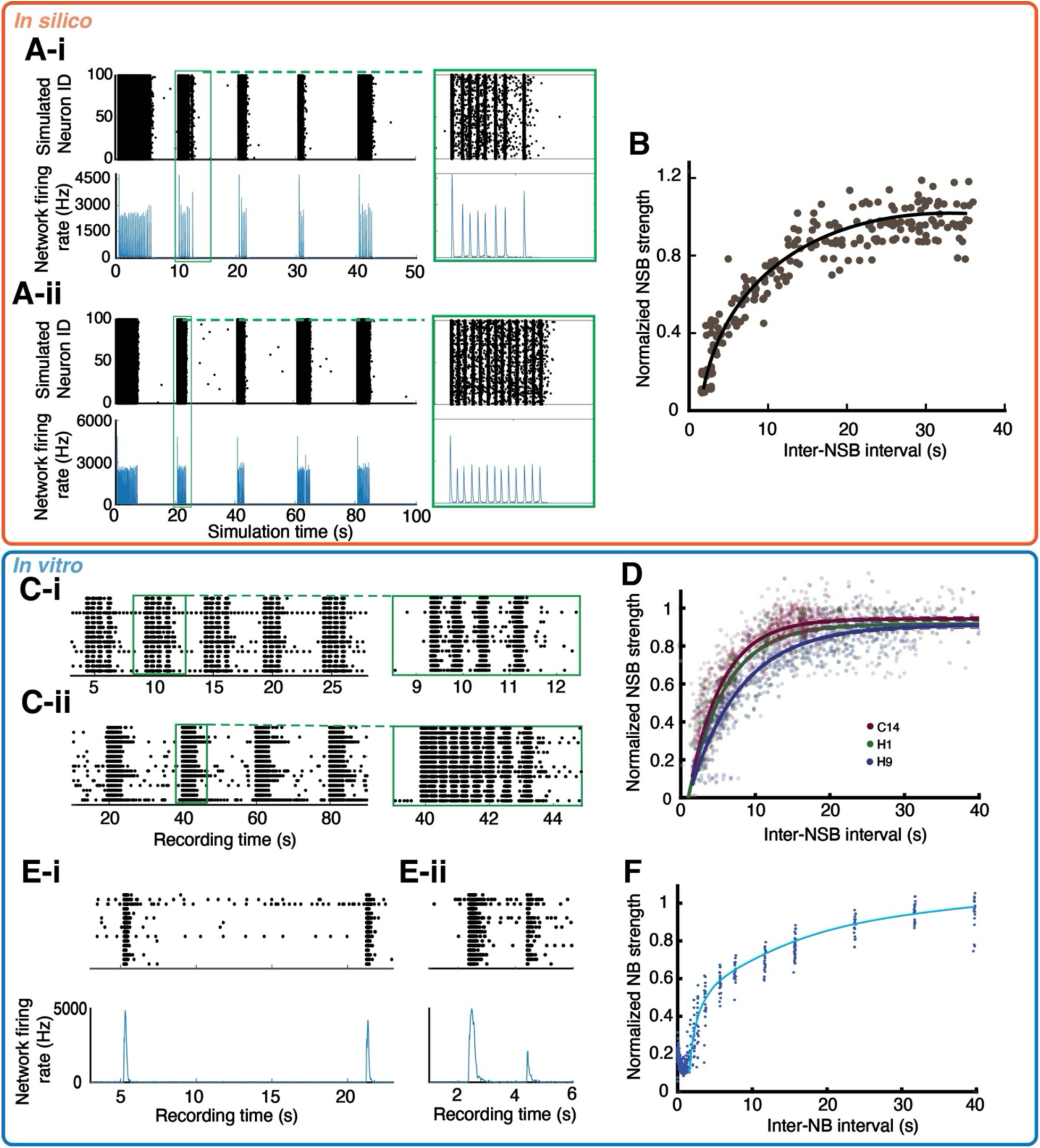
Epileptiform NSBs in iGlutN cultures are associated with presynaptic vesicle pool dynamics. (**A**) Example computational simulations of an excitatory network responding to stimulations with 10s (i) and 20s (ii) intervals. (**B**) Exponential increase of NSB strength (measured by enclosed spike count) in relation to stimulation interval. The fitted curve’s time constant of 9.2s closely matches the model’s 10s RP recovery setting. Data collected from 20 simulations; strength of NSBs normalized to first induced NSB (i.e., when the presynaptic RP is full) in each simulation. (**C**) NSBs elicited at intervals of 5s (i) and 20s (ii) in a 7-week-old culture. (**D**) NSBs show an exponential recovery curve when elicited at different intervals. Within each culture, the strength of elicited NSBs was normalized to the mean strength of spontaneous NSBs. Shown are exponential fits of NSB changes in 3 genetic backgrounds. Recovery time constants are 6.5s, 5.7s, and 8.6s, respectively. (**E**) In a 7-week-old iGlutN culture treated with 5 μM CNQX, two paired pulse electrical stimulations of different intervals elicit single NBs of different strength. (**F**) Strength of the second elicited NBs normalized to the first elicited NBs during paired-pulse stimulations and plotted against the intervals between paired pulses, showing a two-phase recovery curve. Data collected from 7 partially inhibited 7 to 10-week-old iGlutN cultures.

Subsequent *in vitro* experiments were then conducted with iGlutN cultures. A single pulse electrical stimulation could activate the iGlutN networks and elicit an NSB *in vitro*. Activating the cultures with electrical stimulations of larger intervals elicited stronger NSBs (Fig. 2C_ii_ vs. 2C_i_), and plotting the induced NSB strength against inter-NSB interval also yielded an exponential recovery curve with a time constant of approximately 6 seconds, resembling that of RP recovery (Fig. 2D). Additionally, to demonstrate the association between single NBs and presynaptic RRP, we adapted the ‘paired pulse ratio’ paradigm, which is commonly used to study synaptic vesicle dynamics by calculating the ratio of the amplitude of the second induced postsynaptic response to that of the first. When the glutamate receptors were partially inhibited, single NBs but not NSBs could be elicited by electrical stimuli (Fig. 2E). And activating the cultures with paired stimulations of different intervals then yielded a double-exponential recovery curve of single NBs, resembling that of RRP recovery (Fig. 2, E_i_, E_ii_ &F).

### Inhibition of actin polymerization to decelerate RP-to-RRP vesicle translocation modulates NB and NSB dynamics differently

Assuming the slowing presynaptic RRP replenishment terminates the epileptiform NSBs, direct interference of RP-to-RRP vesicle transport should inhibit the development of NSBs. Transportation of presynaptic vesicles from RP to RRP has been shown to be actin-dependent (Miki et al., 2016; Sakaba and Neher, 2003). And inhibition of actin polymerization specifically slows the rapid recovery of RRP and led to a ‘right-shift’ of the double exponential recovery curve (Babu et al., 2020) (Fig. 3A, schematic illustration). We thus applied the actin polymerization inhibitor cytochalasin-D (CytoD), which shortened and terminated NSBs in a dose-dependent manner (Figs. 3B-C). While DMSO did not significantly alter NSB profile (relative strength 1.03 ± 0.08, p = 0.76), 400 nM CytoD significantly reduced NSB strength (relative strength 0.47 ± 0.12, p < 0.001), and 4 μM CytoD further attenuated NSBs to near-single burst level (relative strength 0.11 ± 0.06, p < 0.001). CytoD also led to a dose-dependent ’down-shift’ of the recovery curve of induced NSBs (Fig. 3D). By contrast, in iGlutN cultures partially inhibited by 5 μM CNQX (see Fig. 2E), application of CytoD did not affect the maximum strength of induced NBs but caused a ‘right-shift’ to the NB recovery curve (Figs. 3E-F), resembling the effect of CytoD on RRP recovery (Babu et al., 2020). These wet lab results could also be replicated in our *in silico* model where retarded presynaptic RP-to-RRP vesicle transportation led to a ‘down-shift’ of NSB recovery curve and a ‘right-shift’ of NB recovery curve (Supplementary Fig. S4). Collectively, these findings support the hypothesis that the dynamics of presynaptic vesicle transportation from RP to RRP determines the development of epileptiform NSBs in excitatory neural networks.

**Figure 3.**
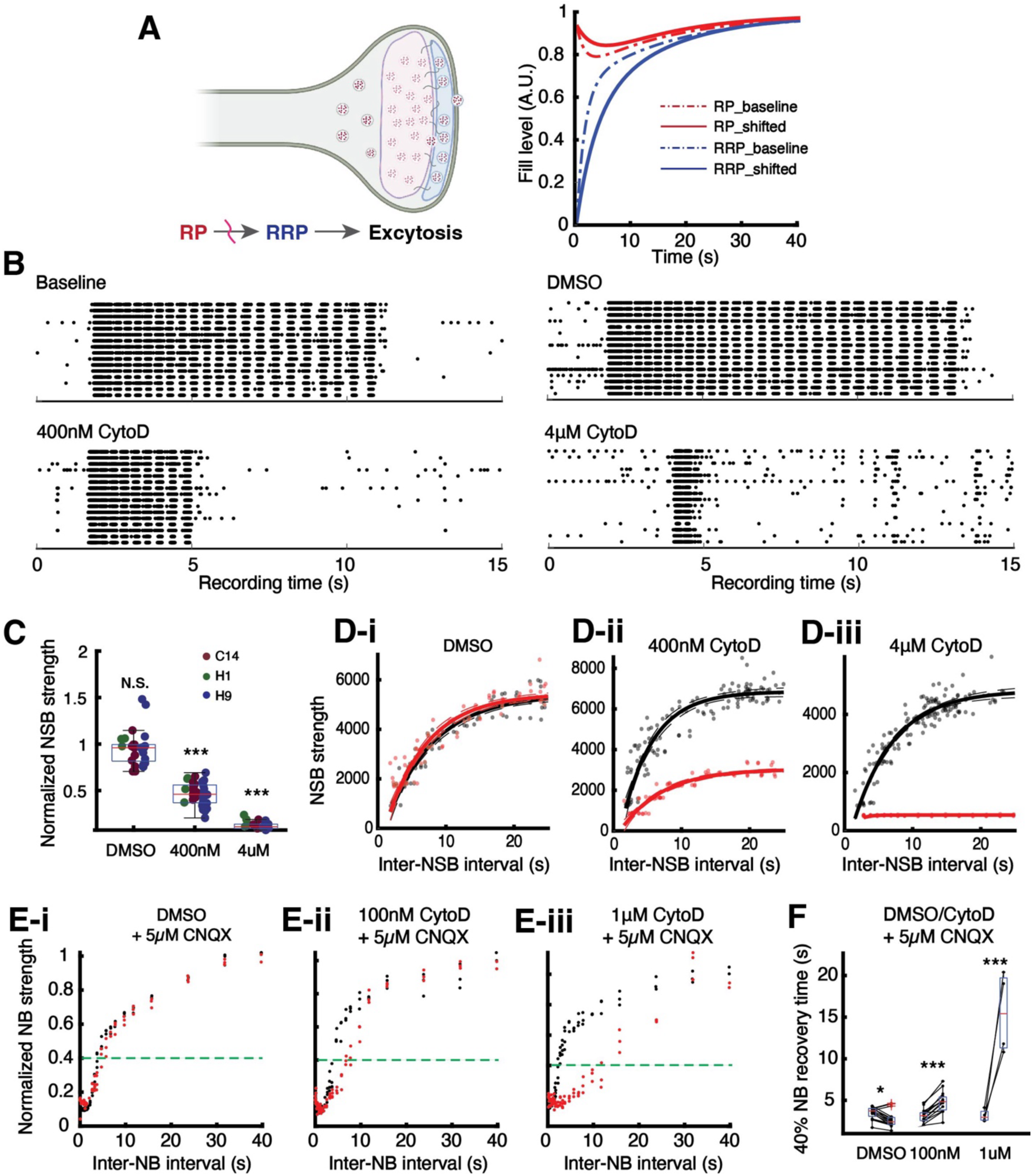
Inhibition of actin polymerization as a tool to attenuate presynaptic RP-to-RRP vesicle transport and to modulate NSBs and NBs. (A) Schematic depiction of the proposed effect of actin polymerization inhibition on the translocation of presynaptic vesicles and the recovery curves of presynaptic RP and RRP. (B) Example raster plot of NSBs generated in the presence of different dosages of CytoD. (C) Change of spontaneous NSB strength upon treatment with 400 nM and 4 µM CytoD. Each point represents one independent MEA culture ( n = 3, 8, and 9 cultures across C14, H1, and H9). (**D**) Example NSB recovery curves (line C14) before (black) and after (red) treatment with DMSO (i), 400nM CytoD (ii), and 4μM CytoD (iii). (**E**) Example NB recovery plots (line C14) before (black) and after (red) application of DMSO (i), 100nM CytoD(ii), and 1μM CytoD(iii) in iGlutN cultures partially inhibited by 5 µM CNQX. (**F**) Change of time to reach 40% NB recovery after treatment with 100 nM and 1 µM CytoD in partially CNQX-inhibited iGlutN cultures (line C14, n = 5 cultures). N.S., not significant; ***, p < 0.001.

### Tonic-clonic transition of epileptiform activities in excitatory neuronal networks mediated by RP-to-RRP vesicle translocation

Upon further exploration of the in silico model, we noticed that with the same parameter settings and initial conditions, simulated networks in different runs would either show regular slowing clonic bursting or start with a tonic phase that transitions into clonic bursting, akin to tonic-clonic seizures (Fig. 4A) — a phenomenon which could also be observed in one of our cell lines (Supplementary Fig. S2B). We reasoned that tonic and clonic activities reflect two closely associated states of excitatory networks rather than a single irreversible progression and thus investigated whether the slowing RP-to-RRP vesicle transport might play a role in eliciting this pattern.

**Figure 4.**
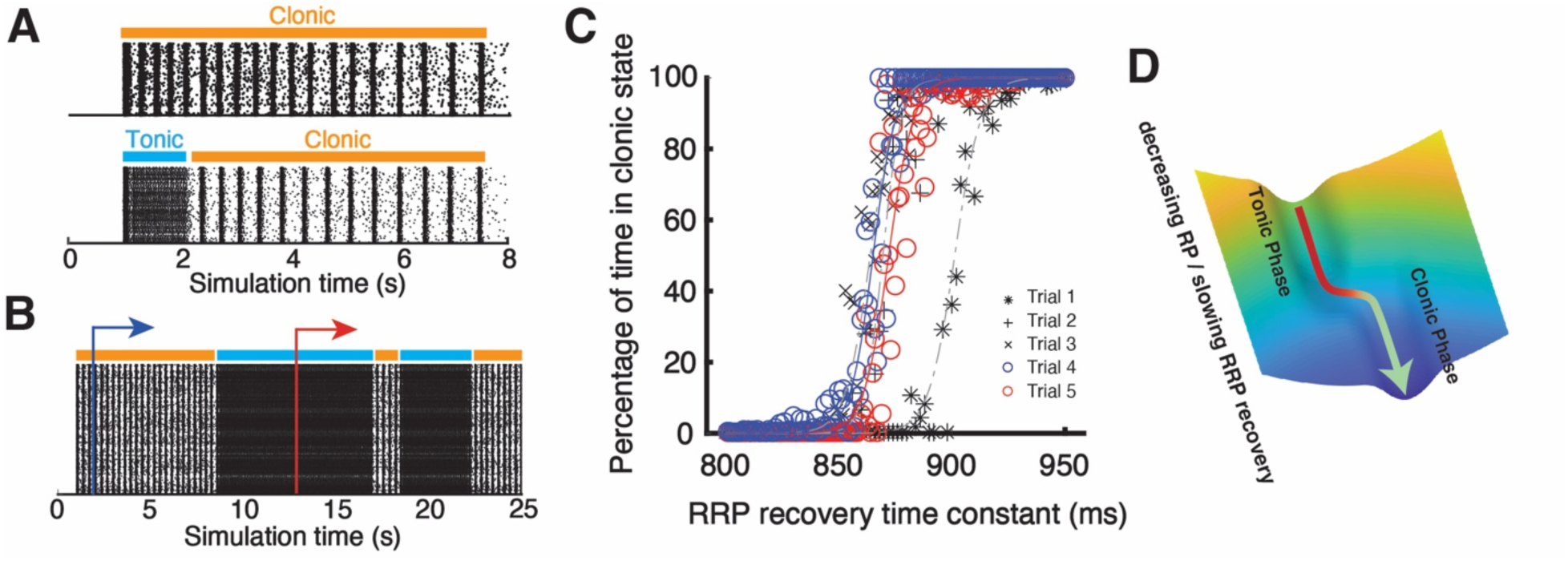
Synaptic vesicle transport from RP to RRP mediates tonic-clonic transition in epileptiform activities. (A) Example raster plots showing a regular slowing clonic NSB (upper panel) and a tonic-to-clonic transition (lower panel) produced by networks with identical parameters in silico. (B) Example in silico raster plot showing bidirectional tonic-clonic switching when RRP recovery was reduced to a single-exponential process (i.e., a constant RP, as in Supplementary Fig. S3). Blue and red arrows mark two time points at which the same trial was branched and continued under different fixed τ*_RRP_* values, corresponding to Trials 4 and 5 in (C). (**C**) Percentage of time in the clonic state in networks with the RRP recovery time constant τ*_RRP_* fixed at different values. Each trial is an independent sweep of the same network; Trials 4 and 5 share an identical history up to their respective branch points in (B) and therefore differ in absolute clonic proportion while preserving the same dependence on τ*_RRP_*. (**D**) Schematic of tonic-to-clonic transitions due to decreasing RP and slowing RRP recovery.

This notion is supported by the observation that when RRP recovery was reduced to a single-exponential process (i.e., a constant RP, as in Supplementary Fig. S3), the network switched repeatedly between tonic and clonic states within a single run (Fig. 4B), indicating the two states are mutually accessible rather than sequential. This bidirectional switching suggests that excitatory networks possess an intrinsic tonic-clonic bi-stability.

We next asked what governs the balance between the two states. Simulations were performed with the RRP recovery time constant (τ*_RRP_*) fixed at different values. With faster RRP recovery the network was predominantly tonic, and as τ*_RRP_* increased the proportion of the run spent in the clonic state rose steeply across a narrow τ*_RRP_* window, until at sufficiently slow recovery time constants only clonic activity persisted (Fig. 4C). While the timing of individual transitions was stochastic, this clonic proportion was reproducibly determined by τ*_RRP_* — even when a single trial was branched at two different time points and continued under different fixed τ*_RRP_* values (blue and red arrows, Fig. 4B; Trials 4 and 5, Fig. 4C), the two branches differed in absolute proportion but preserved the same overall dependence on τ*_RRP_*.

Taken together, these results suggest excitatory networks possess a tonic-clonic bi-stability, with slowing RRP recovery increasing the probability of clonic states (Fig. 4D). Notably, the bidirectional switching in Fig. 4B required holding RP constant; in more realistic settings, where RRP recovery slows progressively rather than being held fixed, the network is biased toward the clonic state from the outset, so predominantly one-way tonic-to-clonic transitions — as in Fig. 4A — would be expected.

## Discussion

In this study, we identified and characterized a nested network super-burst (NSB) phenomenon in human PSC-derived glutamatergic networks that mirrors the rhythmic 4– 2 Hz slowing of EEG discharges seen in human absence seizures and the clonic phase of tonic-clonic seizures. By integrating *in vitro* human neuronal cultures with *in silico* modeling, we demonstrate that the hierarchical organization of presynaptic vesicle pools specifically the transition from the recycling pool (RP) to the readily releasable pool (RRP) — is a primary driver of these complex epileptiform dynamics.

Current understanding of seizure dynamics has largely emphasized GABAergic inhibition as the primary mechanism for generating rhythmic spike-and-wave oscillations (Destexhe and Sejnowski, 1995) and seizure termination (Boido et al., 2014; Curtis and Avoli, 2015). However, the emergence of NSBs in a near-pure excitatory network (Rhee et al., 2019) where GABA receptor antagonists failed to modulate the activity — suggests that intrinsic dynamics of glutamatergic synapses are sufficient to account for complex seizure-like rhythms. The computational model incorporating a hierarchical presynaptic RRP recovery and activity-dependent short-term facilitation successfully recapitulated the observed NSBs. Experiments with electrical stimulation then demonstrated the correlation between the epileptiform NSB and the RP, as well as NB and the RRP as they share comparable recovery time constants (Rizzoli and Betz, 2005; Neher, 2015). Decelerating RP-to-RRP vesicle translocation also differentially modulated NB and NSB in neuronal cultures as predicted *in silico*, indicating that epileptiform dynamics depend heavily on this translocation process.

Glutamate depletion/replenishment and asynchronous release have been proposed to modulate synchronized bursting activity in slice preparations (Staley et al., 1998; Jones et al., 2007). In their earlier work, the authors documented two distinguished time constants of glutamate recovery and bursting strength, with the latter only manifesting after GABA blockage (Jones et al., 2007). They did not, however, point out a sequential or hierarchical relationship of the two processes. It is likely that, in the presence of inhibition, a burst could complete only a single round of RRP depletion and recovery: GABA release counteracts the asynchronous-release-mediated facilitation, preventing subsequent bursts from being induced. Without GABAergic inhibition, a burst or super-burst instead proceeds until the bifurcation point at which facilitation decay outpaces RRP replenishment.

It has been shown that there are distinct steps before presynaptic vesicles can be primed and readied for release (Neher and Brose, 2018; Rizzoli, 2014). These studies were mainly performed on the single cell level with artificial modulation of presynaptic release. How these steps may manifest on a macro-network level and modulate neural activities under physiological conditions has yet to be elucidated. The present work provides a direct link by demonstrating that the hierarchical organization of presynaptic vesicle pools serves as a primary driver of macro-scale network rhythms under physiological conditions. This alignment effectively bridges the gap between fundamental synaptic properties and the emergent complexity of neuronal network activity.

In sum, this study identifies epileptiform network super-bursts (NSBs) as a robust emergent property of highly excitatory human neuronal networks derived from human PSCs via NGN2-mediated forward programming. These findings demonstrate that intrinsic glutamatergic synaptic dynamics alone are sufficient to drive these complex rhythms. Specifically, presynaptic RP-to-RRP vesicle translocation, along with short-term facilitation, appear to play a central role in prolongation and termination of epileptiform activity. Additional simulation work also offered a novel, inhibition-independent explanation for the tonic-to-clonic transition that characterizes generalized seizures: a progressive slowing of presynaptic vesicle replenishment within excitatory networks may itself be sufficient to drive this transition, without requiring changes in inhibitory drive.

Notably, this resonates with the antiseizure efficacy of agents targeting the synaptic vesicle cycle, such as levetiracetam, and suggests presynaptic release dynamics as a mechanistic axis worth further exploration in seizure modulation. From a methodological perspective, the present study illustrates that combining MEA recordings of defined PSC-derived neuronal cultures with computational simulation can provide mechanistic insight into human neuronal network dysfunction.

### Limitations of the study

The results of our study suggest that epileptiform rhythmic activity can emerge from randomly connected glutamatergic neuronal networks and be explained by the hierarchical organization of presynaptic vesicle pools. However, we did not rule out the contribution of other factors in our system, e.g., postsynaptic modulation, changes of ion concentrations inside and outside neurons (Antonio et al., 2016), or the impact of astrocytes which contribute greatly to glutamate recycling (Zhan et al., 2011). Furthermore, the relationship between burst activities and presynaptic vesicle recycling was only indirectly demonstrated. Critically, further refinement of the specificity of presynaptic vesicle modulation is needed as actin polymerization inhibition may have broader biological consequences. In addition, the extent to which the current framework could be applied to in vivo situations remains to be validated. Moreover, while the present study deliberately focused on pure glutamatergic networks to mechanistically isolate the presynaptic excitatory component without inhibitory confounds, NSBs were also observed in bicuculline-treated excitatory-inhibitory mixed cultures (Supplementary Fig. S2D). Further studies are required to explore the validity and generality of the proposed framework, and to explore how inhibitory interneurons modulate NSB dynamics.

## Methods

### In vitro human neuronal cultures

hPSC lines carrying doxycycline inducible NGN2 were established from an induced pluripotent stem cell line generated in-house from skin fibroblasts of a healthy Caucasian male (iLB-C14m-s11; abbreviated C14, registered at hPSCreg as UKBi017-A (https://hpscreg.eu/cell-line/UKBi017-A), and human embryonic stem cell lines WA01 (aka H1) and WA09 (aka H9) obtained from WiCell Research Institute (Wisconsin, USA). Generation and use of iPSC was approved by the Ethics Committee of the University of Bonn Medical Center (approval number: 275/08). The subject gave written informed consent. Work on hESCs and derivatives was approved according to the German Stem Cell Act (Robert-Koch-Institute, Berlin; permit #117). For each parental hPSC background, one previously characterized inducible NGN2-transgenic line was used. Human iGlutNs were produced and cryopreserved at day 8 after doxycycline induction as described (Peitz et al., 2020). For neuronal cultures, 24-well plates with MEAs (M384-tMEA-24 W, Axion BioSystems) were coated with Matrigel (1:30 dilution, Corning, 354230). After thawing, iGlutNs were counted, reseeded in neuronal medium (Neurobasal Medium (Thermo Fisher Scientific, 21103-049), 2% B27 supplement (Thermo Fisher Scientific, 17504-044), 1% GlutaMAX (Thermo Fisher Scientific, 35050-38), and 10 ng/mL BDNF (Cell Guidance Systems, GFH1)) at 1000/mm^2^. Primary mouse astrocytes were added (2000/mm^2^) one day later. The co-cultures were maintained in neuronal medium supplemented with 1 µg/mL doxycycline and 0.5% FBS, with medium change performed twice per week. Culture age is reported in completed weeks after plating cryopreserved iGlutNs.

### MEA recording and electrical stimulation

A Maestro MEA EDGE system and the control software Navigator (Axion BioSystems) were used for signal recording and electrical stimulation. For recording of spontaneous activity, plates were loaded and equilibrated for 20–30 mins, then recorded for 10 mins. Raw data were recorded with the sampling rate at 12.5 kHz, bandpass between 0.1 and 2000 Hz. For pharmacological experiments, compounds were added at ratio of 1:20 to reach a working concentration.

To electrically stimulate the culture, biphasic voltage pulses predefined in the Navigator software (‘Neural Stimulation’ setting, amplitude = 100 mV, duration = 400 μS) were delivered to the target electrodes. Predefined electrical stimulation protocols were used to elicit NSBs in iGlutN cultures to obtain the recovery curve. Specifically, in each session, 5 stimuli with fixed interval between 3s and 30s were delivered to the cultures, with 5-minute-long resting periods between sessions. For the characterization of induced NBs in mature iGlutN cultures, a pre-experiment was performed for each culture to determine the concentration of glutamatergic receptor antagonists such that electrical pulses could robustly elicit single NBs but not propagated epileptiform NSBs. We found 5 μM CNQX usually yielded satisfactory results, but optimization may be needed depending on the cell line and cultivation time. We adapted the ‘paired-pulse paradigm’ widely used in synapse research to better elucidate the parallels between the dynamics of presynaptic RRP and NBs in our cultures. Specifically, paired-pulses with intervals between 0.5s and 40s were applied to the iGlutN cultures, and between paired-pulses were 1-minute-long resting periods. For each culture, the session was repeated 3 times.

### Data processing and statistical analysis

Recorded data were processed with the Navigator software and customized scripts in MATLAB (The Mathworks Inc., R2018b). Online spike detection was performed with a threshold of 6 x SD. The output .spk files containing waveforms and time stamps of detected spikes were then processed offline. NB and NSB detection was performed using an algorithm reported previously (Bakkum et al., 2014). Specifically, a single-channel burst is marked if 5 spikes occurred in less than 100 ms, and a network burst (NB) is marked when 70% of all channels showed burst activity. A network super-burst (NSB) was defined by at least 3 successive NBs with intervals less than 1000 ms. The strength of the NBs and the NSBs were defined by the enclosed spike count. For the characterization of NSB recovery, the first induced NSB in each session was omitted and the recovery time of each NSB was defined as the interval between the ending of the previous NSB and the onset of the present NSB. For the characterization of NB recovery, the strength of the second NB relative to the first NB elicited in each paired pulse was calculated.

Unless otherwise stated, one MEA well/culture was treated as the biological experimental unit; repeated stimulation sessions or repeated pulses within the same culture were treated as technical or repeated-measure observations. Statistical analysis was performed using MATLAB. N-way analysis of variance was used in experiments with nested and repetitive designs, and *post hoc* paired comparisons were then corrected using the Tukey-Kramer method. Custom equation fitting was performed using the nonlinear least squares method provided in the Curve Fitting Toolbox of MATLAB.

### In silico computational simulation of biological neuronal network activity

Computational simulations were conducted using BRIAN2 (Stimberg et al., 2019). We employed exponential leaky integrate-and-fire neuronal models defined as:

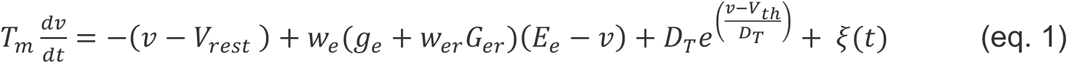

where *T*_m_ is the membrane time constant, *v* is the membrane potential, *V_rest_* is the resting membrane potential. *w_e_* is the adjusting factors of the weight of synaptic input. *g_e_* denotes the conductance of glutamate receptor-coupled sodium channel and has a form of:

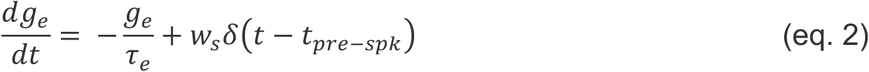

Where *g_e_* increases upon each excitatory presynaptic spike at *t_pre-spk_* and decays with a time constant *τ_e_* . And *w_er_* accounts for the strength of spike-triggered short-term facilitation *G_er_* relative to *g_e_*. And *G_er_* has the form:

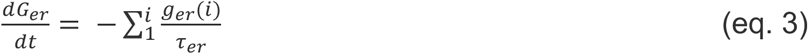

Where *τ_er_* = 800ms. And *g_er_(i)* denotes conductance contributed from the i^th^ synapse this neuron receives. *g_er_(i)* would be updated to 1 upon each presynaptic spike.

*D_T_* and *V_th_* define the exponential threshold where *D_T_* = 0.5 mV. Spikes are defined as *v* >= *V_th_*+ 10 mV. Then *v* would be reset to *V_reset_*. *ξ(t)* is the thermal noise and has a standard deviation of 0.5 mV.

The hierarchy of presynaptic vesicle pools is defined as:

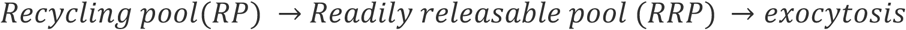

Assuming the upstream vesicle reservoir is infinite, RRP has a volume of 1, RP has a volume of N, then:

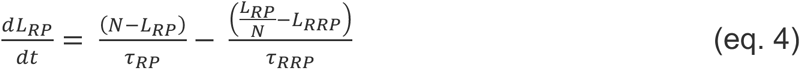

And:

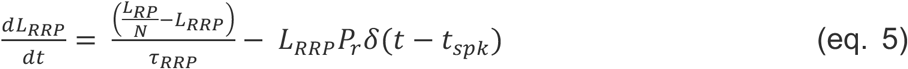

*L_RRP_* serves as *w_s_* in eq. 2.

Values used in the models are listed in Supplementary Table S1. To construct a network, 1000 neurons with the settings described above were connected randomly with a probability of 0.03. To mimic spontaneously firing neurons, a Poisson input unit with 5 Hz mean firing rate was connected to the network. Simulated pulse stimulation was realized by depolarizing targeted neurons by 35 mV.

## Supplemental information

Document S1. Figures S1–S4 and Table S1

## Resource availability

### Lead contact

Requests for further information and resources should be directed to and will be fulfilled by the lead contact, Oliver Brüstle.

### Materials availability

This study did not generate new unique reagents.

### Data and code availability

All data reported in this paper will be shared by the lead contact upon request. All original code has been deposited at Zenodo and is publicly available at https://doi.org/10.5281/zenodo.21053019 as of the date of publication.

## Acknowledgments

This work was supported by stipends to W.J. from the China Scholarship Council (China) and the University of Bonn International Studies Program, which receives support from the Federal Ministry of Education and Research and the Ministry of Culture and Science of the German State of North Rhine-Westphalia under the Excellence Strategy of the Federal and State Governments.

## Author contributions

J.W., M.P., and O.B. conceptualized the study; J.W. and J.L. designed and performed experiments; J.W. and J.L. analyzed the data; J.W. and O.B. wrote the manuscript, with input from all authors; O.B. supervised the work; O.B. acquired financial resources.

