## Supplementary materials for "Presynaptic mechanism of epileptiform activities in forward-programmed human excitatory neuronal networks"

### Supplementary Figure S1

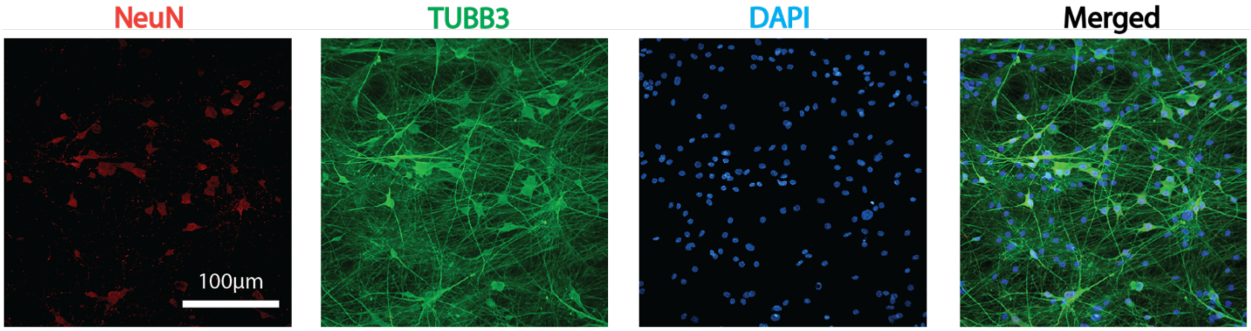


**Supplementary Figure S1.** Representative immunofluorescence images of an iGlutN culture stained for NeuN (red), TUBB3 (green), and DAPI (nuclei, blue), with merged image (right). Scale bar, 100 µm.

Supplementary Figure S2

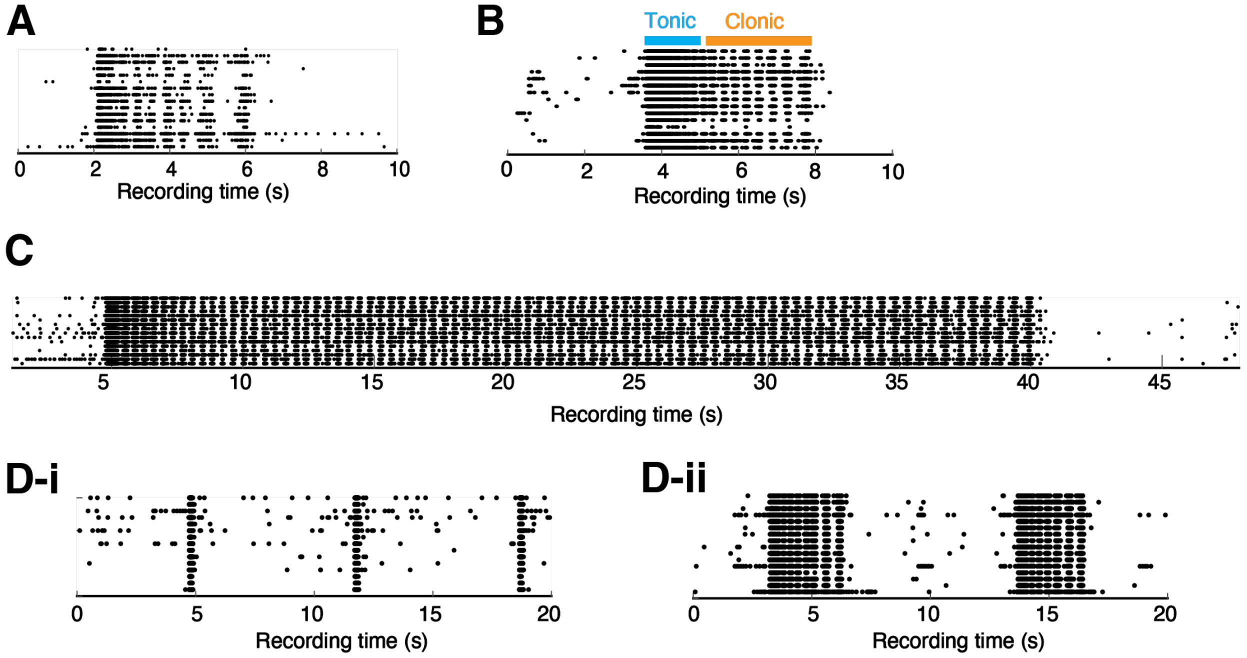


**Supplementary Figure S2.** Variants of epileptiform NSB activities. **(A)** An example ‘immature’ NSB observed in a 4-week-old iGlutN culture. (B) Tonic-clonic-like NSB observed in an iGlutN culture aged 7 weeks (tonic and clonic phases are indicated in blue and orange, respectively). (C) An ultra-long NSB observed in a 7-week-old iGlutN culture. (D) A 7-week-old culture composed of 80% iGlutN and 20% induced GABAergic neurons displayed single network bursts (i) and NSB-like activity after treated with 10uM bicuculline (ii). Induced GABAergic neurons were generated from the C14 iPSC line by doxycycline-induced overexpression of ASCL1 and DLX2 as previously described (Peitz et al., 2020).

Supplementary Figure S3

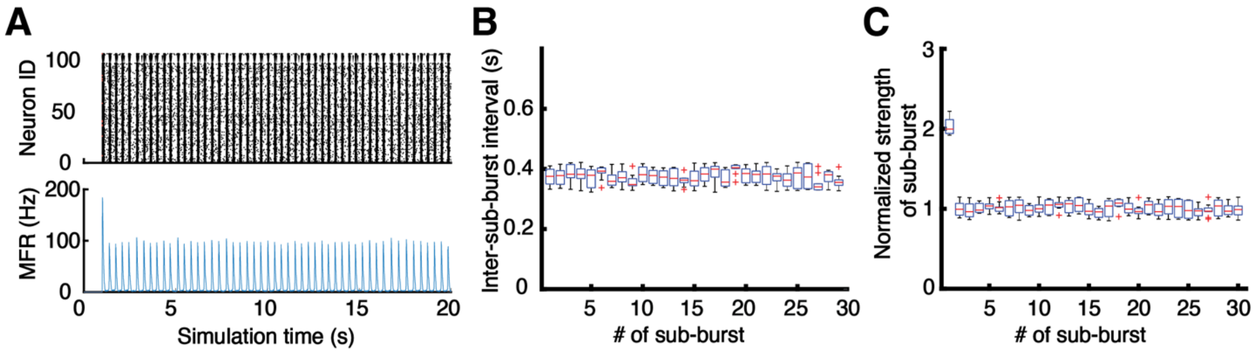


**Supplementary Figure S3.** Unstoppable rhythmic bursting activity in simulated excitatory neuronal network with synaptic short-term facilitation. (A) An example raster plot of a simulation showing unstoppable rhythmic bursting activity. (B-C) Change of IBIs (B) and normalized sub-burst strengths (C) along rhythmic bursting. Data are collected from 20 simulations.

Supplementary Figure S4

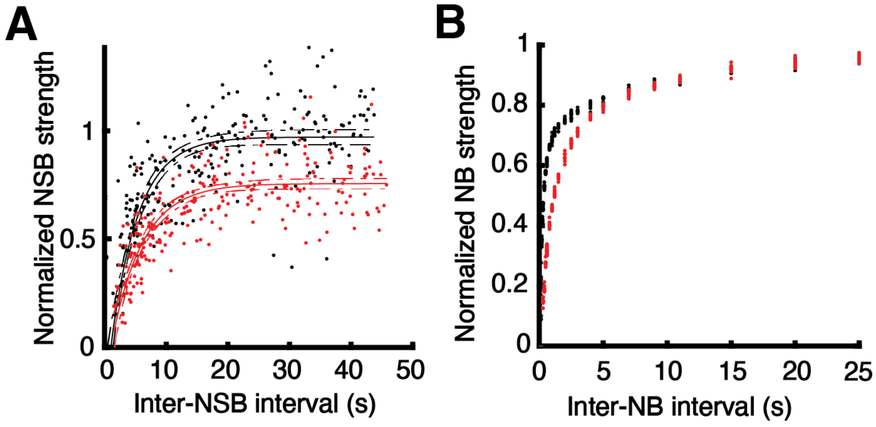


**Supplementary Figure S4.** Change of NSB and NB in simulated network. (A) NSB recovery curve in simulated networks before (black) and after (red) basal rate of RP-to-RRP transportation decreased by 20% (realized by increasing $\tau_{RRP}$ by 20%, see Methods). (B) NB recovery curve in simulated partially inhibited networks (realized by decreasing postsynaptic channel conductance $g_e$ by 30%, see Methods) before (black) and after (red) basal rate of RP-to-RRP transportation decreased by 20%.

Supplementary Table S1. Simulation parameters

| Symbol | Value |
| --- | --- |
| $T_m$ | 38.5 ms |
| $V_{rest}$ | -50 mV |
| $w_e$ | 3 |
| $E_e$ | 0 mV |
| $V_{th}$ | -25 mV |
| $V_{th}$ | -25 mV |
| $\tau_{RRP}$ | 800 ms |
| $P_r$ | 0.25 |
| $V_{reset}$ | -65 mV |
| $w_{er}$ | 0.005 |
| $\tau_{er}$ | 800 ms |
| $\tau_{RP}$ | 7000 ms |
| $N$ | 9 |
